# Pre-movement changes in sensorimotor beta oscillations predict motor adaptation drive

**DOI:** 10.1101/2020.01.13.903807

**Authors:** Henry T Darch, Nadia L Cerminara, Iain D Gilchrist, Richard Apps

**Affiliations:** School of Physiology, Pharmacology and Neuroscience, University of Bristol, BS8 1TD; School of Psychological Science, University of Bristol, BS8 1TU

## Abstract

Beta frequency oscillations in scalp electroencephalography (EEG) recordings over the primary motor cortex have been associated with the preparation and execution of voluntary movements. Here, we test whether changes in beta frequency are related to the preparation of adapted movements in human, and whether such effects generalise to other species (cat). Eleven healthy adult humans performed a joystick visuomotor adaptation task. Beta (15-25Hz) scalp EEG signals recorded over the motor cortex during a pre-movement preparatory phase were, on average, significantly reduced in amplitude during early adaptation trials compared to baseline or late adaptation trials (p=0.01). The changes in beta were not related to measurements of reaction time or duration of the reach. We also recorded LFP activity within the primary motor cortex of three cats during a prism visuomotor adaptation task. Analysis of these signals revealed similar reductions in motor cortical LFP beta frequencies during early adaptation. This effect was also present when controlling for any influence of the reaction time and reaching duration. Overall, the results are consistent with a reduction in pre-movement beta oscillations predicting an increase in adaptive drive in upcoming task performance when motor errors are largest in magnitude and the rate of adaptation is greatest.

## Introduction

Motor adaptation, the ability to modify movements in response to changes in the environment to maintain accuracy, is a fundamental feature of most types of goal-directed behaviour. Motor adaptation is supported by a supraspinal neural network that includes (but is not limited to) the cerebellum and motor cortex. These brain regions are thought to serve differing, but complementary roles during motor adaptation. For example, a large body of work indicates the cerebellum is critically involved in the generation of sensory prediction errors that drive the adaptation process, as well as online control of complex movements ^1–4^. In contrast, the motor cortex appears to support memory of different environment dynamics: Transcranial magnetic disruption of the motor cortex in humans does not influence learning of a new perturbing force, but leads to a deficit in behavioural performance compared with controls in a delayed re-test of the same force^5^, and non-invasive electrical stimulation of the motor cortex can improve retention of motor adaptation performance^6^. Taken together, these data suggest that motor cortical areas support a long-term memory trace of established motor programs, into which newly learned dynamics (potentially computed initially in the cerebellum) are integrated.

Electroencephalography (EEG) signals in the beta frequency band are usually defined as being between 13-30 Hz^7,8^. They are a prominent feature of motor areas in both human and non-human primates and a link between beta frequencies and movement-related processing in the sensorimotor cortex is well-established ^9^. Typically, beta frequency EEG responses in the motor cortex are reduced in amplitude (desynchronised) during movement and increase in amplitude at the end of movements (beta rebound)^8^. Beta amplitude also increases during periods of sustained holding^10–12^ and has been linked to various kinematic parameters including reaction time ^13^ and movement speed^14^.

In the context of motor adaptation, few studies have examined motor cortical beta signals in peri-movement time periods. The beta rebound has been observed to be attenuated during trials in which large errors are made^15^. This is thought to reflect a role in feedback error processing, possibly indicating an adaptive drive (defined here as the period during motor adaptation when motor errors are largest in magnitude and thus the need to modify subsequent motor actions greatest). Conversely, motor cortical beta signals have also been reported to be attenuated prior to movement initiation in both a forcefield and a visuomotor adaptation task^16^. This pre-movement reduction in beta power was interpreted to reflect a predictive updating of the upcoming motor command in response to a previous error, and thereby signals an adaptive drive^16^. However, this idea was challenged by a subsequent study that used an interlimb cooperative adaptation task to decouple adaptive drive from altered motor commands and found that premovement beta modulation might be a result of higher-level processing of sensory afferents that updates upcoming movements^17^. To test the possibility that changes in beta activity in motor cortex could be related to adaptive drive and are generalizable across species we report motor cortical beta frequency activity during motor adaptation in two species: human and cats. We focus on beta activity in the pre-movement period when adaptive drive is greatest for upcoming task performance. This pre-movement period is also not confounded by the execution of the movement (which differs across the two species) allowing a direct comparison of results between the two lines of experiment.

We found that beta frequency power recorded over the motor cortex as scalp EEG in humans, or directly from the primary motor cortex as LFP activity in cats, was reduced (desynchronised) prior to movement in trials when endpoint errors occurred during a joystick visuomotor adaptation task (humans) or when target mis-reaches occurred in a prism visuomotor adaptation task (cats). In both cases the effect was restricted to early adaptation, consistent with a role of reduced beta activity in the motor cortex reflecting adaptive drive.

## Methods

### Ethics

Animal experiments were carried out in accordance with the UK Animals (Scientific Procedures) Act 1986 regulation and were reviewed and approved by the University of Bristol Animal Welfare Ethical Review Body. The human experiments were approved by the University of Bristol Faculty of Science Ethics Committee. All procedures were carried out in accordance with the approved guidelines and. participants provided written informed consent prior to participation. Participants were remunerated for their time in accordance with local policy.

## Behaviour

### Human experiments

Eleven healthy young adults took part (4 female, aged between 21 and 31). All were right-handed (Edinburgh Handedness Inventory score: +87.5, SD±14.3) and had normal or corrected to normal vision.

Participants performed a visuomotor adaptation task requiring them to use a joystick to control an on-screen cursor^15,18^. Each trial was initiated by the participant depressing the joystick trigger. One of four possible black target dots (45°, 135°, 225°, or 315° from vertical) immediately appeared at the edge of an invisible boundary circle centred on the starting position of the user-controlled green dot, forming the task arena (Figure 1). Targets were pseudorandomly presented in blocks of 20 trials, with each target location presented 5 times within each block. Participants were instructed to move the joystick as quickly and accurately as possible to the target, so the movements were effectively ballistic. Trials that exceeded 750ms between target presentation and completion elicited an on-screen prompt to speed up the movements. The joystick was positioned in front of the dominant (right) hand to the right-hand side of the screen, enabling a comfortable grasp whilst removing the hand from the participant’s view whilst engaged in the task. The task was run with Matlab using Psychtoolbox version 3. Task code was modified from that made publicly available by D. Goldschmidt at https://github.com/degoldschmidt/motor-experiments. Figure 1 depicts the appearance of the video screen during an example trial. At the end of each individual trial, the green cursor froze at the point of crossing the boundary of the task space until the participant began the next trial, delivering a stable accuracy marker after the rapid movement.

**Figure 1.**
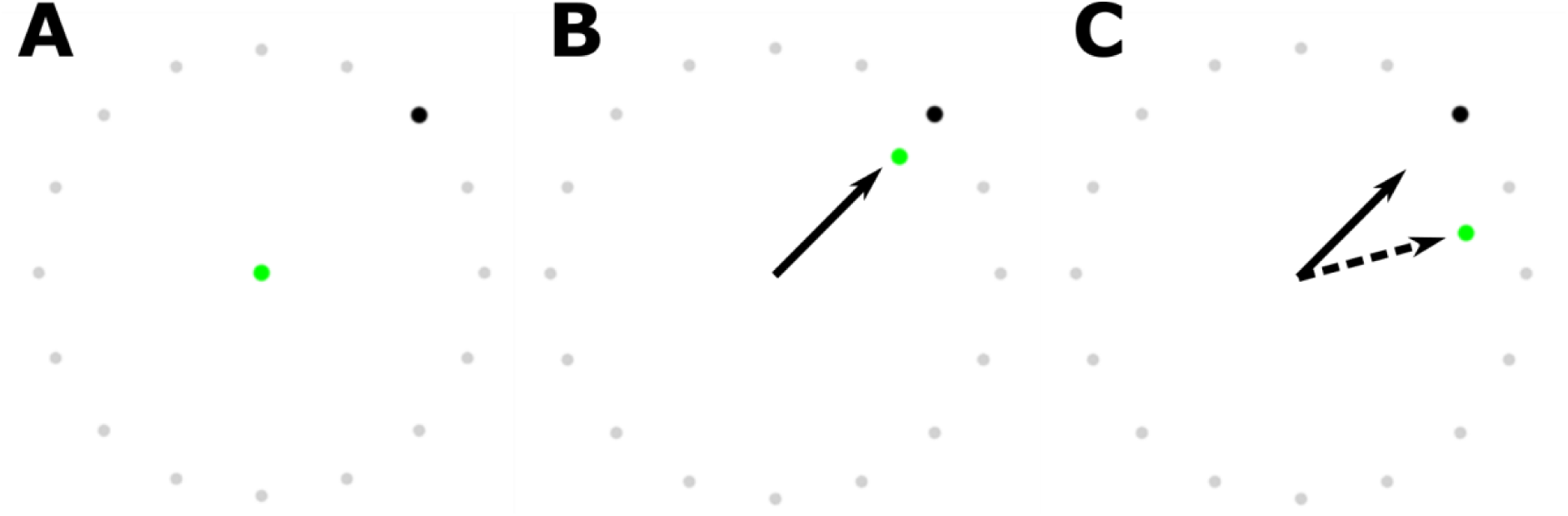
Example view of Human reaching experiment; grey circles indicate the task arena boundary to which participants were instructed to move through. A: View of task arena after clicking the joystick trigger to initiate a trial. One of four black targets and the controlled cursor (green) appear. B: During the first and final trial blocks the green cursor follows the trajectory of the joystick (arrow). C: To induce visuomotor adaptation, a 30 clockwise rotation to the position of the green cursor, providing the illusion of an incorrect joystick trajectory (dashed arrow).

There were three trial blocks each of 200 individual reach trials. The first block consisted of ‘baseline’ trials in which the participants performed the joystick task as normal. The second block induced visuomotor adaptation by implementing a clockwise 35° rotation transformation of the on-screen joystick (green) cursor, requiring participants to modify their outward movement to overcome this perturbation. The final block, identical to the first, was used to probe the participants’ behavioural adaptation to the transformation through the presence of an aftereffect. Participants were not informed of the rotation transformation or the ordering of each block, and none reported explicit awareness of the rotation transform other than that their performance had worsened. Participants did not report the use of an explicit aiming strategy.

Measurements of behavioural error were calculated as the angular displacement relative to the target when reaches crossed the boundary of the task space (endpoint error)^19,20^. Movement initiation was determined when the joystick cursor exceeded a Euclidian distance of 5 pixels from the starting position. Reaction time was measured between the time of target presentation (joystick trigger press), and movement initiation. Reach duration was measured between the time of movement initiation and crossing of the task boundary.

### Animal experiments

Three adult male cats (B, S and G) were trained to perform a left forelimb reach-retrieve task. The target was a Perspex tube (diameter 3cm) placed at eye level at a comfortable reaching height and distance (approximately 20cm) in front of the animal^21^. Reaches were cued by an opaque vertical covering on the Perspex tube which was operated manually by the researchers when the cat was attending to the door and had the reaching limb resting on the ground. The time taken between the door opening and the paw lift-off was considered the reaction time. The reach duration was also recorded as the time between the paw lift-off and entry into the tube, detected by infra-red beams. Reaches were always positively rewarded with a fish morsel within the Perspex tube and no negative training or food restriction was used. Video recordings of the behaviour were made using webcams running at 30fps, synchronised with the neural data acquisition using the open source software Bonsai^22^. These videos were assessed offline to classify reaches (see below).

Once the animals had learnt the reaching task (typically 4 weeks training) surgical implants of recording electrodes were made (see below). Following full recovery from the surgery, daily recording sessions commenced in which the cats performed approximately 80 reaching trials before they became sated and the session was stopped, and they were returned to their home pen. The progression of each recording session followed that of the human study and was divided into three stages. Stage 1: Twenty baseline trials were obtained in which unperturbed reaching was performed and animals rapidly and smoothly placed their forepaw into the target reward tube. Stage 2: Visuomotor adaptation trials in which a set of custom made 40 dioptre Fresnel laterally displacing prism glasses were placed in the animal’s line of sight (approximately 2cm in front of the eyes). Initial trials in the presence of the prism glasses led to reaches laterally missing the entrance of the target reward tube. Adaptation to successfully reach into the target occurred after approximately 10 trials. After 20 consecutive accurate reaches, the prism glasses were removed. Stage 3: immediately after removal of the prism glasses the cats exhibited lateral misses to the opposite side of the tube, indicative of an after-effect.

Based on experimenter observation during the recording session and offline verification by examining a video record of task performance, individual reaches were divided into three categories: (i) ‘Hits’ when the cat rapidly and smoothly reached to place its forepaw into the target reward tube. (ii) ‘Misses’ were the same as hits for the reach component of the movement, but the forepaw failed initially to enter the target, hitting the Perspex façade ∼1 cm to the left or right of the reward tube entrance. A subsequent lateral adjustment was required for the paw to enter the tube to retrieve the reward. (iii) ‘Rejected’ trials were all reaches that either had a reaction time shorter than 150ms; a reach duration (time between paw lift-off and target/façade hit) more than 2 standard deviations of the mean computed for each animal; reaches that displayed a corrective adjustment near the end of the movement, prior to contact with the Perspex façade, in order for the forepaw to enter the reward tube; or trials with large electrical noise artefacts in the neural data. Rejected reaches accounted for approximately 20% of all trials and are not considered further.

### Trial epochs

For the human experiments, the final 20 trials of baseline were used to represent the stable task performance after any initial learning effects at the beginning of the task. The initial 20 trials following implementation of the rotation perturbation were taken to represent early adaptation (EA) in which the adaptive drive (salient error magnitude) is maximal, and the final 20 trials of the perturbation block were taken to represent late adaptation (LA) in which performance is equivalent to baseline (data not shown).

For the cat behaviour, ‘Hits’ made prior to the prism perturbation were taken as equivalent to the baseline condition in the human experiments. The ‘miss’ category is taken as equivalent to the EA period when errors are largest in magnitude and the rate of adaptation is greatest, while ‘Hits’ made during prism perturbation (when adaptation had occurred) were taken as equivalent to the human LA trials.

## Electrophysiological recordings

### Human EEG

The experiments were preformed within a purpose-built Faraday cage. EEG signals were captured using 32 active electrodes (ActiCAP, BrainProducts) arranged according to the extended international 10/20 system. Electrode preparation proceeded as per the manufacturers guidance and impedances were reduced to <5kOhms with conductive gel (SuperVisc, BrainProducts). Data were recorded at 2500Hz using a BrainAmp DC amplifier and BrainVision Recorder Software (BrainProducts).

### Animal LFP

General surgical methods have been reported elsewhere^21^. Briefly, anaesthesia was induced with Propofol (i.v. 10mg/kg, Abbott Animal Health) and maintained with gaseous isofluorane. A veterinary anaesthetist was present to administer anaesthetic agents and monitor vital signs. Stereotaxic surgery involved the creation of a craniotomy over the left motor cortex (approximate coordinates: AP +23, ML +8). An area corresponding to the distal left forelimb motor cortex was identified through microstimulation (11 pulse 330Hz trains^23^ of the cortical surface around the cruciate sulcus. Next, a 16-site silicon probe (NeuroNexus) was implanted into the identified area. The probe consisted of sixteen 2mm shanks, spaced 100µm apart, with a single recording site at each tip. Sites were referenced to a skull screw situated contralateral to the craniotomy. Additionally, EMG recording leads (multistranded stainless steel, Teflon insulated, diameter 0.3 mm, Cooner, Chatsworth, CA, USA) were implanted into the left forelimb Cleidobrachialis Flexor, as well as the long and lateral heads of Triceps Brachii Extensor muscles. A Teflon insulated lead was also implanted into connective tissue in the left forepaw wrist. This lead was used to carry a 400mV 30KHz sine wave transmitted through the paw when in contact with a copper baseplate in the recording environment, enabling the detection of paw lift-off at the initiation of each reach. Animals recovered from surgery in the home pen for one week prior to commencement of daily recording sessions (see above).

Data from the neural probe were captured at 30kHz with a Cerberus neural acquisition processor (Blackrock Microsystems). EMG signals were online filtered with analog filters (bandpass 0.3-5kHz) and visualised on an oscilloscope and then recorded at 30kHz by the Cerberus processor. The paw contact signal was converted into an analog TTL signal to indicate paw contact with the plate and time of paw lift. Paw entry into the tube was determined by a photo-electric switch at the tube entrance.

## Data Pre-processing

### Human EEG

EEG data were offline pre-processed using EEGLab ^24^. For each participant, data were decomposed using the Infomax Independent Component Analysis Algorithm, with components representing noise artefacts (such as eye blinks or neck musculature) identified and removed before back-transforming the data into time-domain electrode signals ^25,26^. As this study was interested in the scalp EEG signals originating from contralateral motor cortex, only data from a single electrode (scalp location - C3) commonly used to represent signals emanating from the motor cortex ^27–30^ was analysed. Trials were identified from the timestamps of the joystick trigger press (target presentation), and movement initiation was detected when the Euclidian distance of the cursor extended beyond a predetermined ‘zero’ zone (5 pixels).

### Animal LFP

Data captured during individual recording sessions were visually inspected in Spike2 (Cambridge Electronic Design) for recording quality and timestamps for reach trials were identified and extracted based on the paw contact signal. LFP recordings across the 16 motor cortical sites appeared highly similar, and no single units were identified on any recording day. Consequently, an example channel was selected, bi-polar referenced to a second channel at the opposite end of the probe (reference site spacing 1.1mm) to remove low frequency interference due to movement.

## Data analysis

Further data analysis was performed using custom made Matlab functions. Pre-processed data from both human and animal experiments were low-pass (100Hz) filtered, down-sampled to 500Hz and high-pass (4Hz) filtered (4th order Butterworth). Next, 2 second windows centred on the movement initiation (joystick movement or paw lift-off event marker) were extracted, and z-score normalised. The time-series were then decomposed into the time-frequency domain using a Morlet wavelet transform (k=6) ^31^, and spectral power computed by taking the log normalised squared magnitude of the wavelet transform. To avoid confounds resulting from smoothing across frequencies as a result of the wavelet transform, the average power in each trial was extracted from a limited range (15-25Hz) within the classical boundaries of beta.

A 200ms analysis window was positioned based on our interest in a period of preparatory activity of data was extracted. In the case of the cats, as we had access to electromyographic (EMG) data from the Cleidobrachialis Flexor, responsible for lifting the forelimb, we were able to determine a suitable location for the window from the trial averaged EMG traces. The flexor muscle is consistently quiescent when the animal is sitting quietly but has a robust ramping activity during reaching that onsets prior to the timestamp of the paw lift (Figure 2). Thus, the 200ms windows were positioned immediately prior to this ramping activity (Cats B and G: 400 to 200ms prior to paw lift. Cat S: 100 to 300ms prior to paw lift). As we did not have access to the equivalent EMG data for human participants, the window was positioned in an equivalent window 400 to 200ms prior to detection of the joystick movement. In all cases, the positioning of the window of interest was such that it fell in between the mean trial initiation and movement initiation events, as such this window represents neural activity occurring during the preparation of the executed reaching movements.

**Figure 2.**
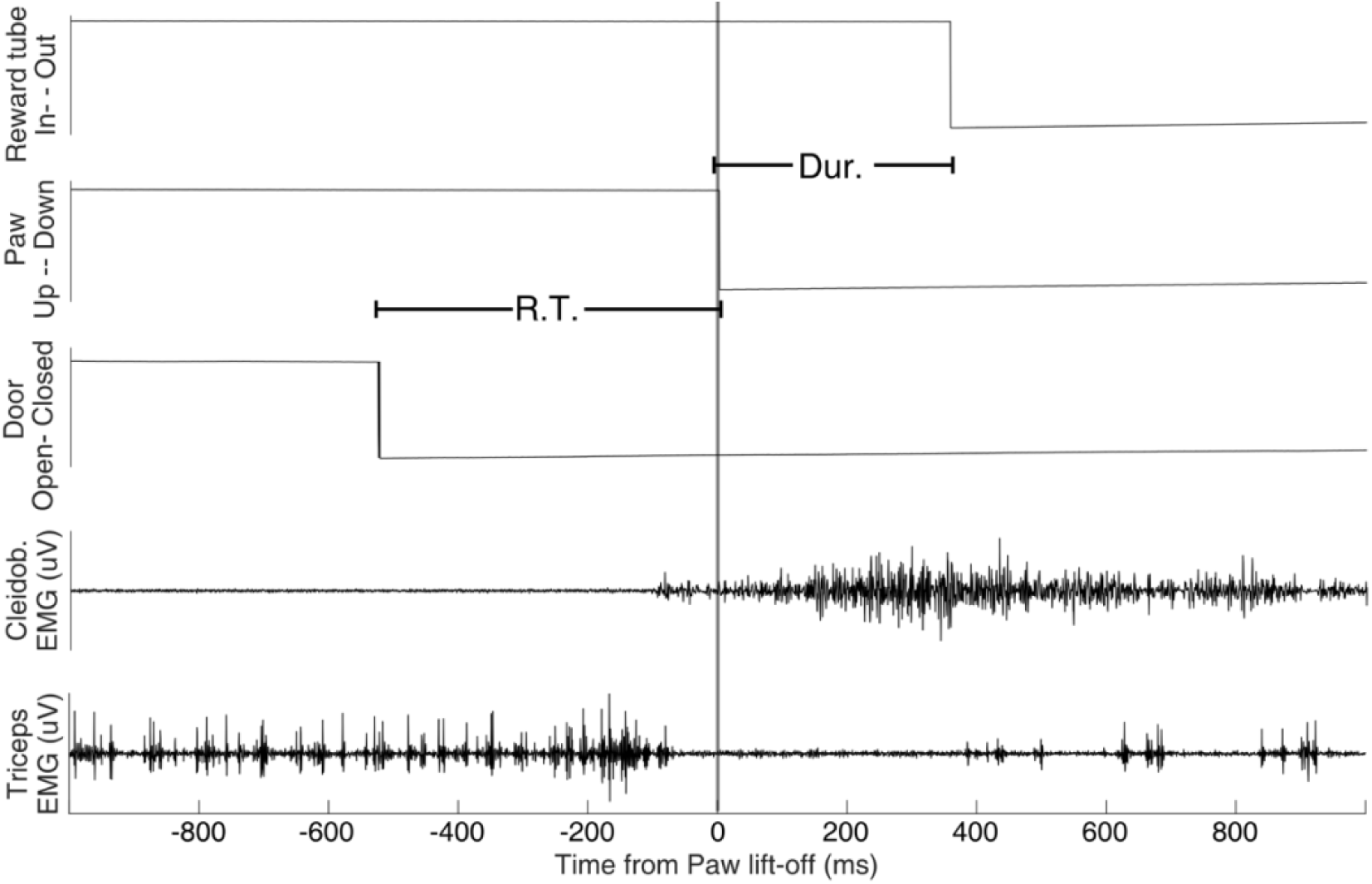
Event markers of a representative trial, with the associated EMG recordings of Triceps (extensor) and Cleidobrachialis (flexor) muscles. Indication of the measured reaction time (R.T.) and duration of the reach (Dur.) from paw lift (time zero) to tube entry are shown.

## Experimental design and statistical analyses

These experiments are formally exploratory as prior to the experiment there was no suitable data to carry out a formal a-priori power calculation. In addition, ethical issues restricted the number of animals in the study. All statistical analyses were performed in SPSS (IBM).

### Human data

Within participants, trials from the same adaptation epoch of interest (Baseline, EA and LA) were averaged to generate a single summary value for each epoch of interest. Data from all participants was then subjected to a repeated measure General linear model (GLM) assessing the effect of adaptation condition, with Bonferroni corrected pairwise post-hoc comparisons. Correlational analyses used Parametric (Pearson’s) methods after visually inspecting data for approximate normality using P-P and Q-Q plots.

### Animal data

Each animal was analysed as an individual case study. Data from each animal were subject to a Univariate ANOVA assessing the effects of adaptation condition. Reaction times and durations were incorporated as covariates. Post-hoc pairwise comparisons were made using Bonferroni correction. Correlational analyses used Parametric (Pearson’s) methods. For all tests, data were checked for normality and homogeneity of variance where appropriate.

## Results

### Human Data

Figure 3 summarises the results in the human joystick adaptation task. A repeated measures GLM with within-subjects comparisons revealed a significant effect of adaptation (F(2,20)=9.10, p=0.002). Beta power in early adaptation (EA) was reduced on average by 0.12dB (37%) by comparison to the baseline condition and reduced on average by 0.16dB (43%) by comparison to late adaptation (LA). These differences were statistically significant (pairwise comparison EA vs Baseline p=0.038 and EA vs LA p=0.010). The small increase (0.04dB, 10%) in beta power between LA and baseline was not statistically significant (p=0.816).

**Figure 3.**
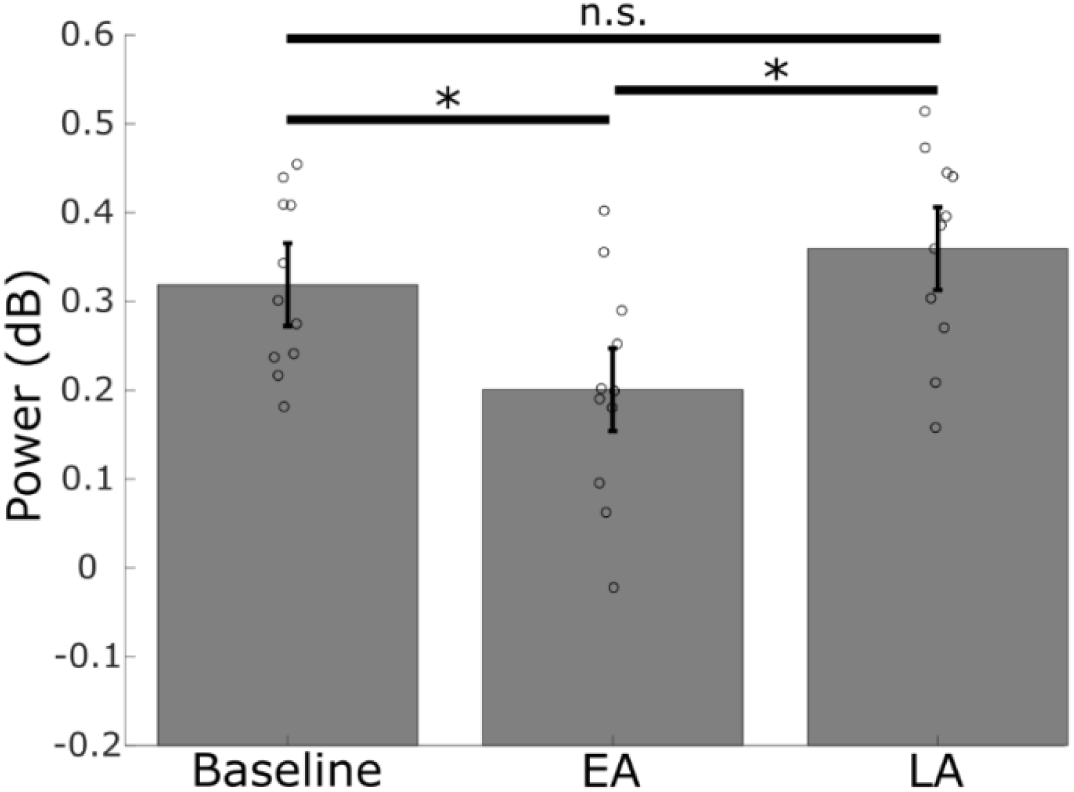
Mean pre-movement Beta (15-25Hz) power recorded over the sensorimotor cortex in human subjects in baseline, early adaptation (EA) and late adaptation (LA). Error bars denote a repeated measures confidence interval, based on α=0.05 ^48,49^. Data for individual participants shown as circles. Asterisks denote statistically significant pairwise contrasts after Bonferroni correction *p<0.05; n.s., p>0.05.

To test whether the differences in beta oscillations could be related to changes in either reaction time or reach duration, the mean change between baseline and EA of beta power, reaction time and duration of the reach was computed for each participant (not shown). This data showed no significant correlation between power and reaction time (Pearson’s r = 0.023, p=0.95) nor duration of the reach (Pearson’s r = -0.048, p=0.89).

### Animal Data

Figure 4 shows equivalent data for the prism adaptation task in cats. In cat S (Figure 4A) the data closely resemble the pooled results from the human study. On average, beta power reduced by 0.445dB in EA compared to baseline and increased by 0.421dB in LA compared to EA. Cat G (Figure 4B) displayed a similar pattern insofar as beta power was reduced in EA compared to both baseline and LA. On average, beta power in EA was decreased by 0.153dB and 0.125dB relative to baseline and LA respectively. Finally, in Cat B (Figure 4C), beta power increased by 0.168dB in EA compared to baseline and decreased by 0.231dB in LA compared to EA.

**Figure 4.**
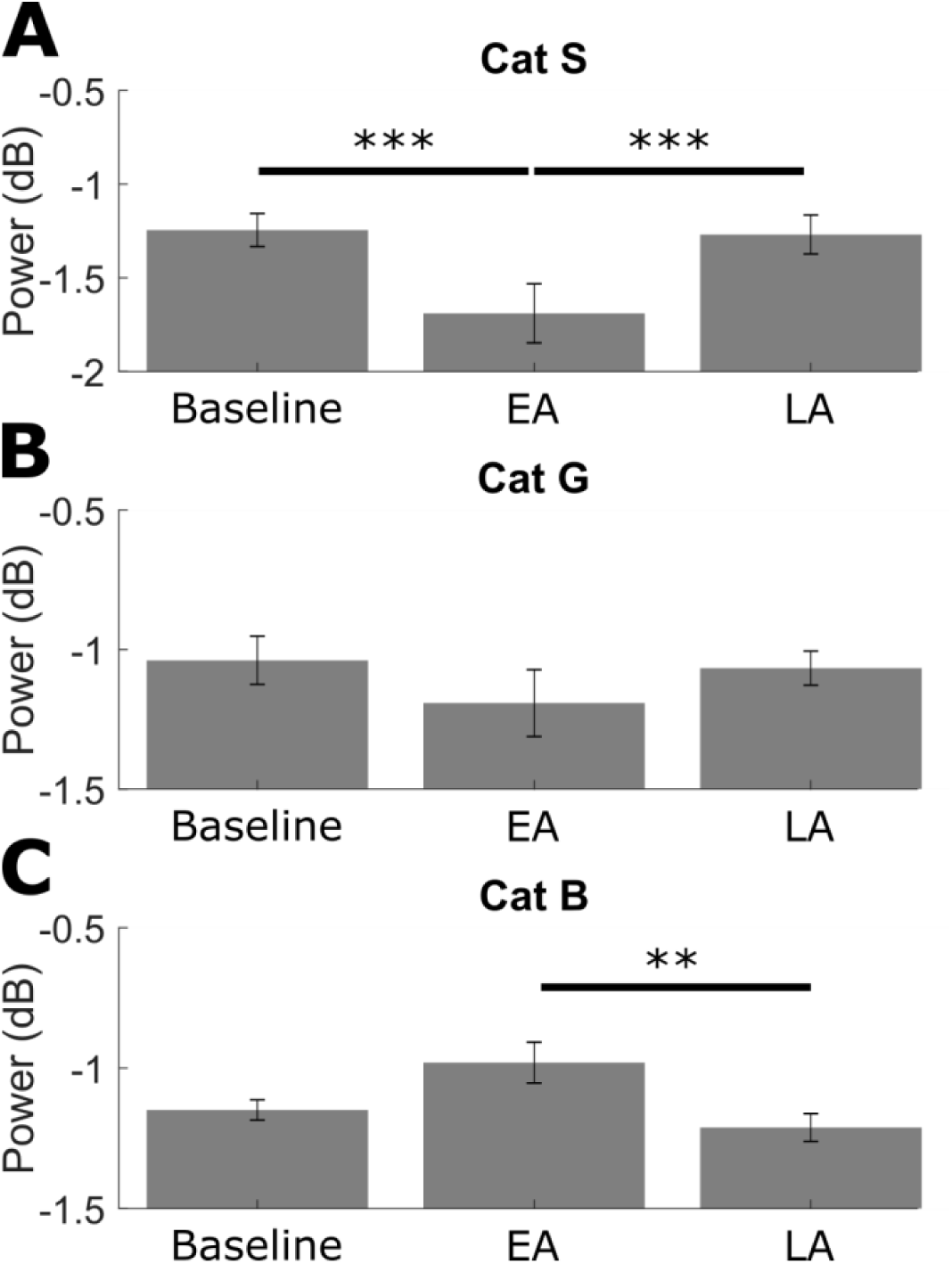
Bar charts of pre-movement beta power from trials in each epoch of interest. Cats ‘S’ and ‘G’ show a decrease in pre-movement beta power during early adaptation trials, increasing again during late adaptation trials (Panels A and B), whilst cat ‘B’ shows an increase in pre-movement beta power during early adaptation trials and a decrease in power during late adaptation trials (panel C). Error bars represent 95% confidence intervals. Asterisks denote significant pairwise contrasts after Bonferroni correction (p<0.0005).

As shown in Table 1, analysis of the LFP beta signal with both the reaction time and reach duration showed a significant correlation for both metrics in one of the three cats (Cat S). In order to account for this potential confound, we included the reaching metrics as covariates in further testing of all animals.

**Table 1.** Statistical summary values of parametric correlational testing between motor cortical beta LFP and two movement parameters: reaction time and duration of the reach.

| CAT ID | METRIC | PEARSON'S R | SIGNIFICANCE<br>(P-VALUE) |
| --- | --- | --- | --- |
| **S** | **Reaction time** | -0.192 | 0.001 |
|  | **Reach duration** | -0.188 | 0.001 |
| **B** | **Reaction time** | -0.014 | 0.784 |
|  | **Reach duration** | -0.046 | 0.367 |
| **G** | **Reaction time** | 0.059 | 0.258 |
|  | **Reach duration** | 0.026 | 0.617 |

Univariate ANOVA (with reaction time and reach duration set as covariates) revealed a significant effect of adaptation condition in cat S (F(2,301)=11.13, p<0.0005), and these differences were statistically significant (pairwise comparisons; Baseline vs EA p<0.0005, EA vs LA p<0.0005). No significant difference was found between baseline and LA (0.024dB increase; pairwise comparisons, p=1.0). The same model for cat G showed no statistically significant effect of the adaptation condition (F(2,365)=1.23, p=0.293). Finally, the Univarate ANOVA model for cat B revealed a significant effect of adaptation condition (F(2,385)=5.430, p=0.005), with significant decrease in beta power between and EA and LA (p=0.006). The increase in beta between Baseline and EA was not significant (p=0.329), nor was there significant changes between Baseline and LA (p=0.057).

In summary, the pooled human EEG data show a reduction in beta power pre-movement in EA compared to both baseline and LA trials. By comparison, in two cats motor cortical LFP exhibited a similar pattern of modulation to the human data, with a pre-movement decrease in EA compared to baseline and LA trials. The exception was one animal that showed a non-significant increase in beta power in EA by comparison to both baseline and LA. These effects were not related to the reaction time or movement duration.

In Figure 5 the cat and human data are compared directly. For the human data, boxplots are shown for the difference in beta power between conditions (error bars indicate maximum and minimum differences). The equivalent data (mean difference between conditions) for the three cats are overlaid as individual data points. The variance and magnitude of effects are comparable between the two species, with the animal LFP data falling mainly within the distribution of the larger sample of human data.

**Figure 5.**
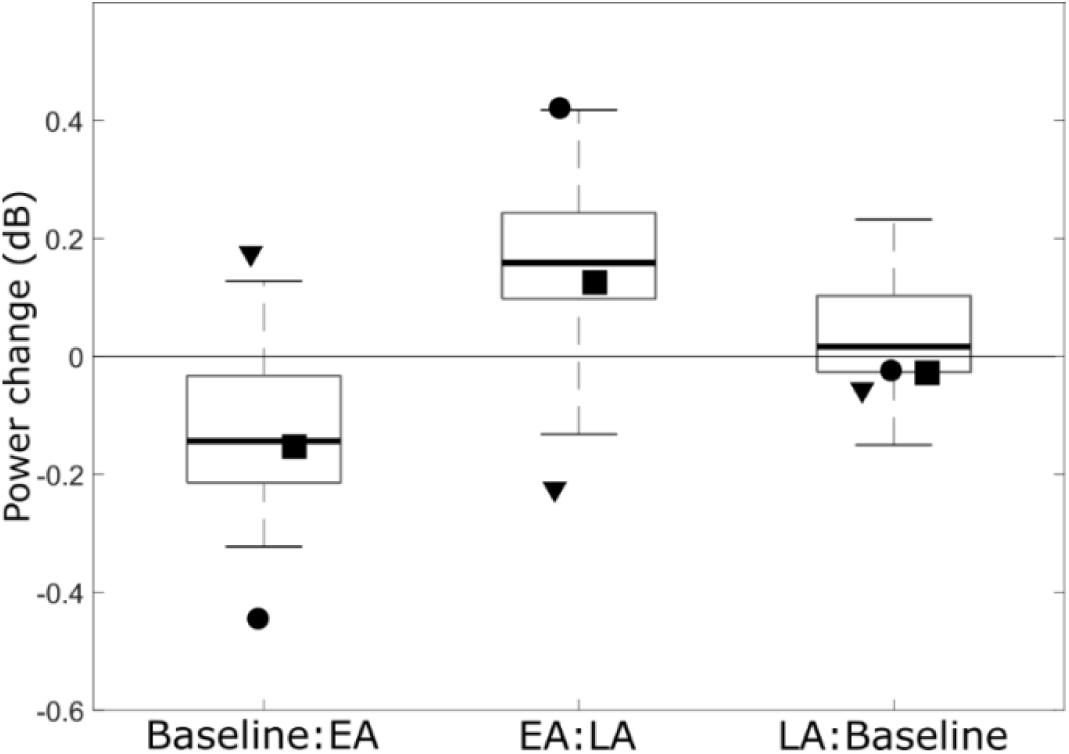
Comparison of effects between humans and cats. Boxplots display contrasts between adaptation conditions for the pooled human data. Boxes indicate interquartile range (central bar median), error bars indicate maximum and minimum range of data. Average data from individual animals shown by markers: • cat S, ▪ cat G, ▾ cat B.

## Discussion

In this study we show that EEG beta frequencies, recorded at a scalp location classically associated with motor cortical signals, are attenuated immediately prior to movement initiation specifically during early stages of visuo-motor adaptation, and returned to baseline levels when behavioural performance (endpoint errors) had returned to baseline performance: i.e. when adaptation drive is greatest. Further, we provide evidence that beta activity recoded as local field potential activity directly from the motor cortex in cats exhibits similar motor adaptation related changes.

Our results are consistent with previous work in human subjects which show a decrease in pre-movement sensorimotor EEG beta power during early stages of forcefield, and visuomotor adaptation task – involving perturbing forces being applied to the limb^16,17^. This would suggest the activity within the motor cortex is independent of the mode of sensory feedback. Forcefield adaptation normally involves the integration of visual and proprioceptive information^32^, whereas visuomotor adaptation relies on visual feedback to monitor performance^33^. This contrasts with previous studies that indicate that forcefield and visuomotor adaptation may involve different neural pathways/processes within the cerebellar cortex ^34–37^. One possible explanation for this difference is that signals arising from distinct cerebellar cortical regions are integrated within the cerebellar nuclei and/or thalamus before being forwarded to the motor cortex for the preparation of subsequent limb movements.

Our results from three cats performing a prism visuomotor adaptation task provide additional novel evidence that a similar pre-movement modulation in beta power occurs in motor cortical LFP signals. Individual differences observed within the animal data were mainly within or close to the expected inter-individual variance observed for the human data. We take this as evidence to suggest that the pre-movement beta effects are generalizable across species, and that the effects seen in the EEG data are at least in part driven by motor cortical activity.

Importantly, the reduction in pre-movement beta activity appeared to be independent of reaction time, which suggests that it is not related to movement initiation, whereas enhanced beta activity has been implicated in a delay of movement onset in both monkeys and humans^13,14^. Pre-movement beta was also not correlated to the duration of reaches, whereas a study of Parkinsonian patients concluded that beta oscillations in the supplementary motor area do appear related to movement velocity^38^. Here, pre-movement beta attenuation was specific to periods where the drive to adaptively modify the upcoming movement was greatest.

Simultaneous EEG, LFP and multi-unit spiking activity recording in area F5 (ventral pre-motor cortex) of macaques have previously shown an inverse relationship between multi-unit activity and LFP and EEG beta power^39^. Therefore, the present pre-movement beta modulation could reflect an increase in motor cortical spiking activity at times of increased adaptative drive. Indeed, beta oscillations have been linked with antidromic activity within the Pyramidal Tract neurons of the motor cortex^40^, as well as the balance of inhibition/excitation (with elevated levels of endogenous GABA enhancing both baseline beta oscillations and the magnitude of the movement-related desynchronisation) ^41–43^. Furthermore, studies of non-human primates during motor tasks has revealed that many cortical neurons synchronise their activity to beta rhythms during periods of oscillation, limiting their spiking rate to a narrower range than that present outside of oscillatory episodes ^44^. Together, this raises the possibility that the pre-movement beta desynchronisation observed here and in previous studies reflects altered inhibitory control during times of increased adaptive drive.

An alternative (but not mutually exclusive) view of the origins of LFP oscillations is that they reflect synchronous activity of synaptic inputs to neurons local to the recording site ^45–47^. The observed modulation of beta power could then be interpreted to represent changes in the amount of neural synchrony in afferent signals to the motor cortex as proposed by Torrecillos and colleagues ^16^.

In conclusion, our results show an adaptation related attenuation of beta oscillations during movement preparation, recorded by scalp EEG over the motor cortex in humans and as motor cortical LFP signals in cats. Our findings are consistent with the reduction in beta power reflecting desynchrony of motor cortical PTNs in the early stages of adaptation when there are large movement errors. This could correspond to changes in the excitatory/inhibitory balance disrupting the ongoing beta oscillations, thus enabling modifications to established motor programs in response to error signals detected by other brain regions such as the cerebellum. Further work is necessary to elucidate the precise mechanisms supporting this beta phenomenon.

## Acknowledgements

This work was supported by a BBSRC funded SWBio DTP studentship, and MRC grant G1100626. The authors are grateful for the support and assistance of Dr Jo Murrell (veterinary anaesthetist) and R. Bissett during the animal experiments, and to Dr N. Kazanina for the use of the high-density EEG lab and equipment.

## Data Availability Statement

The data included in this report are available from the corresponding authors upon reasonable request

## Author Contributions

N.L.C., I.D.G. and R.A conceived and designed the study with input from H.T.D., H.T.D, N.L.C and R.A acquired the data, H.T.D conducted all analyses, all authors contributed to the interpretation of data, drafted the manuscript and agreed with the submitted version or the paper.

## Conflicts of interest statement

The authors declare no competing interests.

